# The nature of intraspecific genome size variation in taxonomically complex eyebrights

**DOI:** 10.1101/2021.04.27.441637

**Authors:** Hannes Becher, Robyn F. Powell, Max R. Brown, Chris Metherell, Jaume Pellicer, Ilia J. Leitch, Alex D. Twyford

## Abstract

- Genome size (GS) is a key trait related to morphology, life history, and evolvability. Although GS is, by definition, affected by presence/absence variants (PAVs), which are ubiquitous in population sequencing studies, GS is often treated as an intrinsic property of a species. Here, we studied intra- and interspecific GS variation in taxonomically complex British eyebrights (*Euphrasia*).
- We generated GS data for 192 individuals of diploid and tetraploid *Euphrasia* and analysed GS variation in relation to ploidy, taxonomy, population affiliation, and geography. We further compared the genomic repeat content of 30 samples.
- We found considerable genuine intraspecific GS variation, and observed isolation-by-distance for GS in outcrossing diploids. Tetraploid *Euphrasia* showed contrasting patterns, with GS increasing with latitude in outcrossing *Euphrasia arctica*, but little GS variation in the highly selfing *Euphrasia micrantha*. Interspecific differences in GS genomic repeat percentages were small.
- We show the utility of treating GS as the outcome of polygenic variation. Like other types of genetic variation, such as single nucleotide polymorphisms, GS variation may be increased through hybridisation and population subdivision. In addition to selection on associated traits, GS is predicted to be affected indirectly by selection due to pleiotropy of the underlying PAVs.

## INTRODUCTION

Genome size (GS), defined as the amount of DNA in an individual’s unreplicated haploid nucleus (Greilhuber *et al*., 2005), is associated with an organisms life history strategy, development, physiology, ecology, and gene and genome dynamics and evolution (Van’t Hof & Sparrow, 1963; Beaulieu *et al*., 2008; Šímová & Herben, 2012; Greilhuber & Leitch, 2013; Simonin & Roddy, 2018; Bilinski *et al*., 2018; Novák *et al*., 2020a; Roddy *et al*., 2020). Genome size is estimated to show a c. 64,000-fold variation across Eukaryotes, and c. 2440-fold variation in flowering plants (Pellicer *et al*., 2018). Much is known about broad-scale variation in GS across land plants and algae, with different phyla characterised by different GS ranges (Pellicer & Leitch, 2020), and showing, in many cases, a strong phylogenetic signal (e.g. Weiss-Schneeweiss *et al*., 2006; Vallès *et al*., 2013; Wang *et al*., 2016; Bainard *et al*., 2019; Cacho *et al*., 2021). Genome size has also been explored in polyploid species, with studies showing that while whole genome duplication events initially lead to an increase in GS, their subsequent evolution is often accompanied by genome downsizing over time (Leitch *et al*., 2008; Leitch & Leitch, 2008; Pellicer *et al*., 2010; Wong & Murray, 2012; Wendel, 2015; Zenil-Ferguson *et al*., 2016). Recently, community ecology studies have started to include data on GS and demonstrate its influence in shaping plant diversity (Guignard *et al*., 2016, 2019). While representative GS estimates have been obtained for approximately two thirds of flowering plant families (Pellicer & Leitch, 2020), variation between individuals and populations within species has typically received less attention, despite the increasing realisation that such variation may be common (e.g. Šmarda *et al*., 2010; Kolář *et al*., 2017).

GS has often been considered a property of a species, and there has been much debate as to whether it genuinely varies within species (Greilhuber, 2005; Gregory & Johnston, 2008; Šmarda & Bureš, 2010). Genuine intraspecific differences in DNA content have been reported or are predicted between individuals with: (1) heteromorphic sex chromosomes (Costich *et al*., 1991; Renner *et al*., 2017), (2) different numbers of B chromosomes (Leitch *et al*., 2007) dysploidy and aneuploidy, or (3) the presence/absence of specific DNA sequences such as (a) structural variants including insertion-deletion polymorphisms (indels), (b) copy number variation in protein-coding genes, commonly found in pan-genome studies (Hirsch *et al*., 2014; Wang *et al*., 2018b; Gao *et al*., 2019; Hübner *et al*., 2019; Göktay *et al*., 2020), and (c) copy number variation of rDNA copies (Long *et al*., 2013) or of other genomic repeats (Chia *et al*., 2012; Haberer *et al*., 2020). Some differences, such as small indels, can be as small as one base pair, while others are large-scale (many megabases), including sequence duplications or loss of a dispensable chromosome. The above types of genetic variation can be subsumed under the term presence/absence variants (PAVs), a type of structural genomic variation, and may be detectable by methods for estimating GS, such as flow cytometry. Modern protocols using flow cytometry with appropriate reference standards, and following best practice approaches, can be accurate and highly precise (Greilhuber *et al*., 2007; Pellicer *et al*., 2021) and reveal genuine intraspecific variation that can be confirmed by genome sequencing. Such sequencing has also been used to reveal that repeat differences can be useful genetic markers, including microsatellites and AFLPs. Consequently, there are an increasing number of well-documented reports of genuine intraspecific GS variation (e.g. Achigan-Dako *et al*., 2008; Šmarda *et al*., 2010; Díez *et al*., 2013; Hanušová *et al*., 2014; Blommaert, 2020).

Our study considers such variation as polygenic, meaning heritable, and with a value affected by multiple independent loci in the genome (Figure **1**). Many polygenic traits are known, with the most renowned example being human height (Fisher, 1919). Loci underpinning polygenic variation need not be protein-coding genes, but may also involve non-coding sequences including introns, promotors, trans elements, or genomic repeats. Loci underpinning a polygenic trait may differ in their effect sizes, as shown by Koornneef *et al*. (1991) for flowering time in *Arabidopsis thaliana* (see also Napp-Zinn, 1955). Further, variants at a genetic locus are commonly pleiotropic, affecting multiple traits and thus potentially being the target of multiple selective effects. An early example of treating GS as such is the study of Meagher *et al*. (2005) on the relationship between GS and flower size in *Silene latifolia*, that showed correlations between floral traits and GS in male plants.

**Figure 1.**
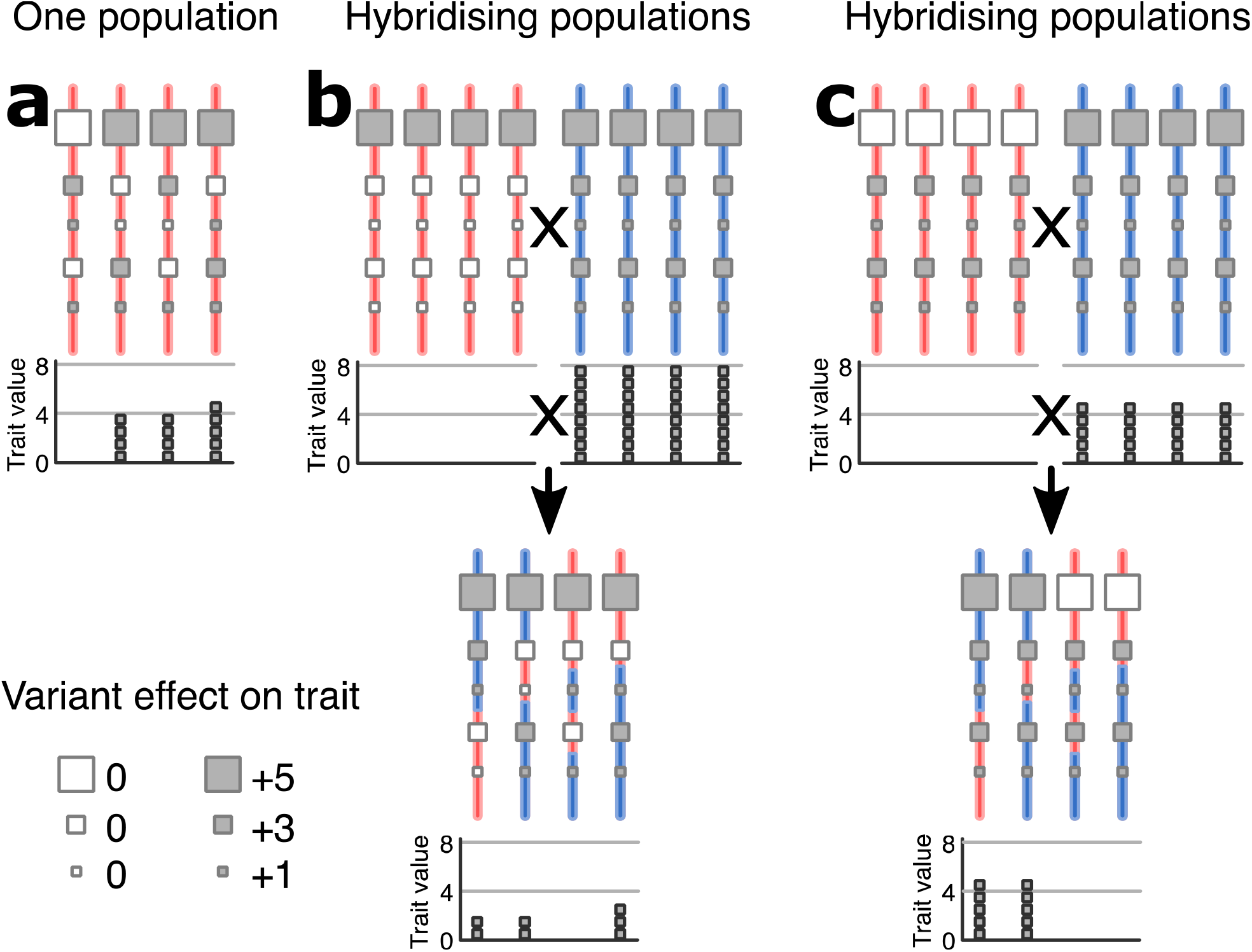
Schematic illustration of a polygenic trait, and its variability after hybridisation. Each red or blue line represents an individual’s genome. Squares represent genetic variants with different effect sizes on a trait. The bar charts indicate individuals’ trait values, relative to the individual with the lowest value. (a) A population (or species) with genetic variability for the trait. The effect of hybridisation between populations with different trait values depends on the genetic architecture of the trait difference. If the populations differ in many variants with small effects (b), recombinant offspring (denoted by mixed red and blue lines) are likely to have similar trait values. If, however, trait differences are due to a few variants with large effects (c), segregation in the recombinant offspring can produce higher trait variation. Applied to GS, open squares correspond to DNA missing and filled squares to DNA present at some site in the genome, as detailed in the main text.

Here we explore GS variation in British eyebrights (*Euphrasia* L., Orobanchaceae), a recently radiating group of facultative hemiparasites. They comprise five diploid (2n = 2x = 22) and 15 tetraploid species (2n = 4x = 44) (Metherell & Rumsey, 2018), with genomic sequencing showing that British tetraploids are closely related allotetraploids, with one sub-genome derived from, or closely related to, British diploids (Becher *et al*., 2020). The genus is an ideal model group for investigating GS variation within and between closely related species because species diversification is frequently postglacial (Gussarova *et al*., 2008; Wang *et al*., 2018a), with many taxa being narrow endemics or recent hybrid species. *Euphrasia* therefore provides multiple opportunities to study GS changes at the early stages of species divergence. Heterogeneous ecological conditions may promote local adaptation, and extensive hybridisation may result in local geographic homogenisation with variation in GS structured by geography rather than by taxonomy, as seen previously in microsatellite and AFLP studies of population structure (Kolseth & Lönn, 2005; French *et al*., 2008).

To investigate the nature of GS variation in British *Euphrasia* species, we generated a comprehensive dataset of 192 GS estimates across 13 species and 10 hybrid combinations, supplemented with genomic sequence data to estimate the abundance of genomic repeats for 30 diverse diploids and tetraploids. Our study aims to answer the following questions: (1) How variable is GS within species, between species, and between ploidy levels? (2) What is the contribution of genomic repeats to GS variation in British *Euphrasia*, and how does repeat content differ between the ploidy levels? (3) Does GS variation correspond with known patterns of genetic structure and/or environmental variables in British *Euphrasia*? We discuss our results in the light of polygenic variation, and we argue for a closer integration of population genomics research with research on GS variation.

## METHODS

### Population and species-level genome size variation

#### Population sampling

Our sampling for GS estimation aimed to collect from across the diversity of British *Euphrasia* taxa, and from a wide geographic area. Samples from 90 populations comprising 13 species and 10 hybrid combinations were either wild-collected and used directly for GS estimates (54 samples) or collected as seeds and grown at the Royal Botanic Garden Edinburgh following Brown *et al*. (2020) prior to GS estimation (138 samples). A full list of samples analysed including their origin is given in Supplementary Information Table S1. The identification of species and hybrids were made by the *Euphrasia* taxonomic expert Chris Metherell, based on morphology.

#### Genome size measurements

Nuclear DNA content of *Euphrasia* samples was estimated by flow cytometry using propidium iodide (PI) stained nuclei, following the one step method (see Pellicer *et al*., 2021). Briefly, for each *Euphrasia* accession, two small leaves (c. 1-2 cm) were chopped together with the internal standard *Oryza sativa* ‘IR36’ (1C = 0.5 pg; Bennett & Smith, 1991) using a new razor blade, in a petri dish containing 1 mL of ‘general purpose isolation buffer’ (GPB; Loureiro *et al*., 2007), supplemented with 3% PVP-40 and 0.4 μL of β-mercaptoethanol. An additional 1 mL of buffer was added to the homogenate, and then this was filtered through a 30 μm nylon mesh to discard debris. Finally, the sample was stained with 100 μl of PI (1 mg/mL, Sigma) and incubated for 20 min on ice. For each accession analysed, one sample was prepared, and this was run three times on the flow cytometer. The nuclear DNA content of each sample run was estimated by recording at least 5,000 particles (c.1,000 nuclei per fluorescence peak) using a Cyflow SL3 flow cytometer (Sysmex-Partec GmbH, Munster, Germany) fitted with a 100-mW green solid-state laser (Cobolt Samba). Resulting output histograms were analysed using the FlowMax software (v. 2.9, Sysmex-Partec GmbH) for statistical calculations. We report only GS estimates for samples where the coefficients of variation (CV) of the sample and standard peaks in the flow histogram were less than 5% (see Supporting Information Figure S1a and b for illustrative histograms of each ploidy level).

Where differences in GS were detected within a species, combined samples containing at least two accessions were prepared following the same procedure as for individual runs. Genuine intraspecific variation was confirmed where multiple fluorescence peaks were identified from the combined run.

Throughout the paper we give 1C values in pg, where necessary converting published GS values reported in Gbp to pg using a conversion factor of 0.978 following Doležel *et al*. (2003).

### Repeat content variation

#### Sequence data generation

We used a combination of existing and newly generated genomic sequencing data to investigate repeat variation in 31 samples comprising seven diploids and 23 tetraploids of *Euphrasia* plus *Bartsia alpina* as an outgroup. We downloaded short-read Illumina data from the sequence read archive (SRA, see Supplementary Information Table S2). These included 18 samples in total, including 12 tetraploid samples from the isolated island of Fair Isle (Shetland, Scotland) generated for the study of Becher *et al*. (2020), which allowed us to study genomic repeat profiles in sympatric populations. This dataset also included a total of six representative diploid and tetraploid species from elsewhere in Britain.

We supplemented this previous data with newly generated sequence data from eleven additional UK samples representing a wider range of species and geographic locations, including 11 UK *Euphrasia* samples, an Austrian sample of *Euphrasia cuspidata* intended as a close outgroup to UK species, and *Bartsia alpina* as an outgroup to the full sample set. Genomic DNA was extracted from 12 silica-dried samples and herbarium material of *E. cuspidata* using the Qiagen Plant Mini Kit (Qiagen, Manchester, UK), and used to prepare NEBUltra PCR-based libraries. Pooled libraries were sent to Edinburgh Genomics where they were run on a single lane of HiSeq 2500 using high output mode with 125 bp paired-end sequencing.

#### Repeat content

We ran the RepeatExplorer2 (RE) pipeline (https://repeatexplorer-elixir.cerit-sc.cz/; Novák *et al*., 2010, 2013, 2020) on a data set of 25,000 randomly selected read pairs of each of the 31 samples (1,550,000 reads in total). This slightly exceeded the maximal number of reads that can be analysed with default settings (which depends on the data). Our dataset was therefore down-sampled to approximately 20,500 read pairs per sample. In comparative RE analyses, read numbers are often supplied in proportion to genome sizes to assure repeats of similar genome proportion can be detected in all samples (e.g. Novák *et al*., 2020a). This logic does not apply here, where the British samples comprise 23 closely related tetraploids and six closely related diploids, with the diploid genome very similar to one of the tetraploid sub-genomes (Becher *et al*., 2020). No matter what genome proportion is chosen per sample, there will always be more of the shared sub-genome than of the sub-genome restricted to tetraploids. To minimise mate overlaps of short insert sizes, each read was trimmed to 100 nucleotides. Further, we only used reads where at least 90 nucleotides had phred quality scores > 30. To analyse the genomic repeat content, we excluded clusters annotated by RE as plastid DNA or Illumina process controls. Our numbers thus deviate slightly from RE’s automatic annotation.

#### Statistical analyses

Most GS analyses were conducted across all individuals or populations. However, for *E. arctica, E. anglica*, and *E. micrantha*, where sampling covered most of their large geographical range in Britain, we also analysed data from each species separately. All analyses were done using R version 3.6.1 (R Core Team, 2019). For analyses of variance (ANOVAs) we used the function aov(). To test whether sample means of GS were significantly different, we used the function t.test(), with Bonferroni correction in cases of multiple testing. To analyse how GS variation was partitioned by ploidy, taxon, and population we used ANOVA. To test the effect of ‘species’, we then re-ran the ANOVAs without hybrids (Table **1**). To test the significance of GS variance differences between species pairs, we divided the population mean genome sizes by each species’ grand mean (centring) and then applied an *F* test (R function var.test()).

We tested the association between GS and latitude using a mixed effect model (R package nlme, function lme()). For species analysed separately, we used linear models. We carried out Mantel tests to assess the relationship between geographic distance and GS difference across all samples as done by Duchoslav *et al*. (2013). Unlike genetic data, which require population information, these Mantel tests could be carried out on individual-based genome size differences or population means. Isolation by distance was assessed using Mantel tests (R package vegan version 2.5-6) with 999 permutations.

To analyse genomic repeat patterns, we used hierarchical clustering and PCA on a matrix of the per-sample genome proportions of the 100 largest repeat clusters in R using the functions hclust() and prcomp(). *Bartsia alpina* was removed from the final PCA data set, because its divergence from *Euphrasia* accounted for most of the variance in the data set, obscuring variation within *Euphrasia*. To identify repeat clusters with large contributions to the first principal component, we selected those clusters which had absolute values > 0.1 in the first eigenvector. We further used binomial-family generalised linear models to estimate the average genomic proportion individually for each repeat cluster. For each estimate, we computed the residual sum of squares as a measure of the variation in genomic abundance between individuals. We used linear models to assess the differences in relative abundance of individual repeat types between ploidy levels.

To investigate a possible association of individual repeat clusters with GS, we used nine tetraploid samples for which we had both an estimate of the population average GS and repeat data (samples marked with asterisks in Supporting Information Table S2). We used the function cor.test() to assess the significance level of any associations between the genome proportion of each individual repeat cluster and population average GS.

## Results

### Population and species-level genome size variation

Genome size estimates from all 192 individuals passed our quality checks. These samples came from 13 different species and 10 hybrid combinations, including 40 diploid and 152 tetraploid individuals (Supporting Information Table **S1**). Our samples covered a particularly wide geographic range for the large-flowered species *E. anglica* (diploid, 552 km) and *E. arctica* (tetraploid, 1152 km), and the small-flowered and highly selfing *E. micrantha* (tetraploid, 962 km).

The mean GS across all tetraploids was 1.18 pg (s.e. 0.004 pg), which is 11% less than twice the mean GS of the diploids (0.66 pg, s.e. 0.008 pg). In the diploids, individual values ranged 1.2-fold, from 0.60 pg in *E. anglica* (population BED) to 0.73 pg in *E. anglica* in Dumfriesshire (E4E0085). In tetraploids there was 1.3-fold variation, from 0.99 pg in *E. foulaensis* in Fair Isle (FIA105) to 1.33 pg in *E. arctica* in Orkney (E4E0033).

Intraspecific GS ranges were widest in *E. arctica* (n = 43) and *E. foulaensis* (n = 13) (both 1.3-fold), and *E. anglica* (n = 23) (1.2-fold). *Euphrasia confusa* (n = 6), *E. nemorosa* (n = 22), *E. pseudokerneri* (n = 9), and *E. rostkoviana* (n = 9) had GS ranges greater than 1.1-fold. While individuals with different GS values were often found in distant populations, such as in *E. anglica* (0.6 pg and 0.73 pg, 525 km apart), and in *E. arctica* (1.04 pg and 1.33 pg, 903 km apart), we also found considerable GS variation between populations in close proximity in *E. foulaensis* (0.99 pg and 1.25 pg, 2.5 km apart) and in *E. confusa* (1.14 pg and 1.32 pg, same population). In all cases, tests to distinguish genuine intraspecific variation from technical artefacts confirmed the GS differences reported between individuals (see Methods and Supporting Information Figure S**1c** and **d**). Generally, we found wider GS ranges in taxa with more populations sampled. A notable exception was *E. micrantha* (GS range 1.14-1.21 pg from 17 individuals analysed from 9 populations, up to 962 km apart), which is discussed below.

In ANOVAs, the vast majority of the overall GS variation was explained by ‘ploidy’, while ‘taxon’ and ‘population’ accounted for smaller significant fractions (Table **1**). ‘Population’ accounted for considerably more variation than ‘taxon’ – 3 or 8 times, depending on whether hybrids were included in the analysis or not. This difference is due to the few data available for most hybrids (Figure **2a**, Supplementary Information Table **S1**). The fact that ‘taxon’ generally accounts for only a small amount of variance is reflected by the near-continuous distribution of GSs within each ploidy level (Figure **2b**). The distribution of tetraploid GS values has two gaps, caused by a few exceptional individuals with extreme outliers in their GS values. While most tetraploid GS values are between 1.07 and 1.26 pg (red horizontal lines in Figure **2b**), six samples had lower (*E. arctica, E. foulaensis*, and *E. foulaensis* x *marshallii*), and seven higher, GS (*E. arctica*).

**Table 1.**
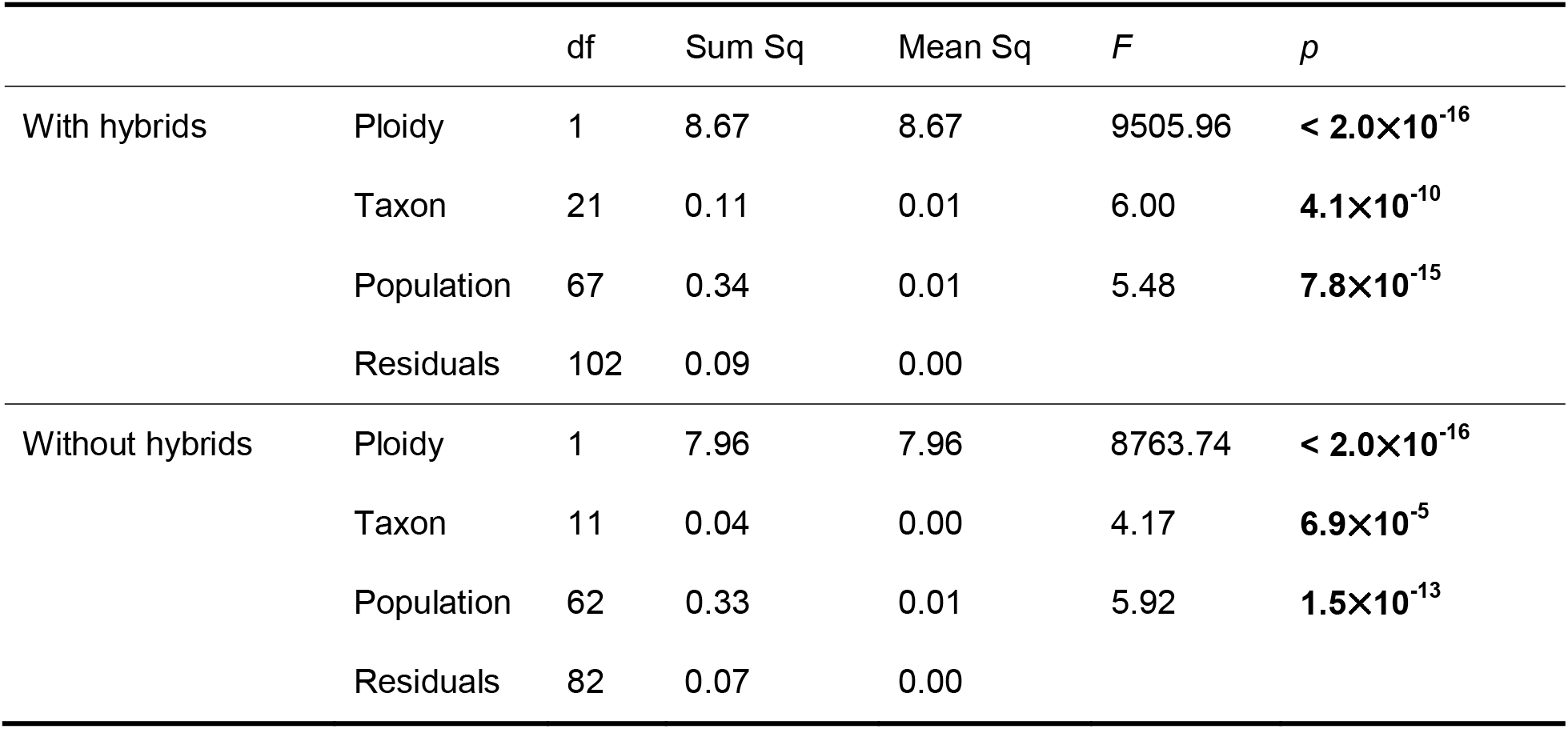
Partitioning of GS variation across *Euphrasia* species and hybrids. Top analysis includes all 192 samples from 13 species and 10 hybrids, and the lower analysis 157 samples comprising just the 13 species (lower analysis). Both ANOVA tables detail the variance components (Sum Sq) accounted for by ploidy, taxon and population.

|  |  | df | Sum Sq | Mean Sq | *F* | *p* |
| --- | --- | --- | --- | --- | --- | --- |
| With hybrids | Ploidy | 1 | 8.67 | 8.67 | 9505.96 | **< 2.0×10<sup>-16</sup>** |
|  | Taxon | 21 | 0.11 | 0.01 | 6.00 | **4.1×10<sup>-10</sup>** |
|  | Population | 67 | 0.34 | 0.01 | 5.48 | **7.8×10<sup>-15</sup>** |
|  | Residuals | 102 | 0.09 | 0.00 |  |  |
| Without hybrids | Ploidy | 1 | 7.96 | 7.96 | 8763.74 | **< 2.0×10<sup>-16</sup>** |
|  | Taxon | 11 | 0.04 | 0.00 | 4.17 | **6.9×10<sup>-5</sup>** |
|  | Population | 62 | 0.33 | 0.01 | 5.92 | **1.5×10<sup>-13</sup>** |
|  | Residuals | 82 | 0.07 | 0.00 |  |  |

**Figure 2.**
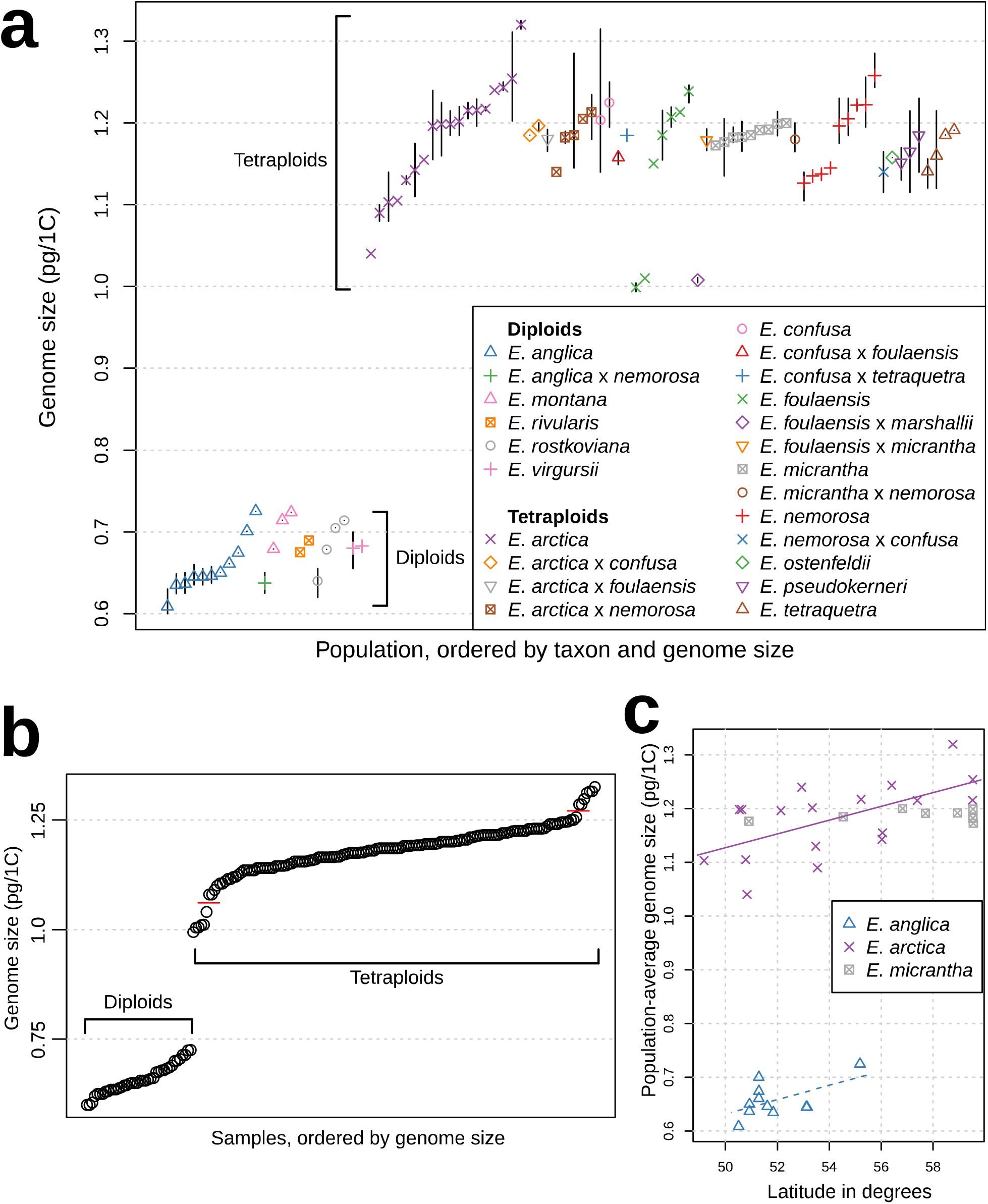
Patterns of GS variation in British *Euphrasia*. **a** The distribution of population-average GS for 90 populations of 23 taxa (13 species and 10 hybrids). Vertical bars indicate the GS range within each population where more than one individual was analysed. **b** Distribution of individual GS estimates for all 192 samples. Horizontal red lines indicate the limits of the continuous part of the tetraploid GS distribution. **c** Population average genome sizes plotted against latitude for the three most widely sampled species. The solid purple line indicates a significant statistical relationship of GS with latitude across 17 populations of *E. arctica*. This relationship was only marginally significant for 11 populations of *E. anglica* (dashed blue line). No significant association was found across nine populations of the highly selfing *E. micrantha*.

Analyses of the three geographically widespread species with wider population sampling revealed that GS variation was significantly partitioned by population for mixed-mating *E. anglica* (*F*_10,12_=9.86, *p*=2.3_×_10^−4^) and *E. arctica* (*F*_17,25_=10.5, *p* < 1.7×10^−7^), but not for highly selfing *E. micrantha* (*F*_8,8_=0.31, *p*=0.94). Further, the variance in population average GS was significantly lower in *E. micrantha* than in *E. anglica* (*F*_10,8_=11.65, *p*=9.6×10^−4^) or *E. arctica* (*F*_17,8_=53. 2, *p*=2.3×10^−6^).

Individual-based Mantel tests to link geographic distance and GS variation were significant over all 40 diploid samples (Mantel statistic *r*=0.25, *p*=0.001) and all 152 tetraploids (*r*=0.04, *p*=0.01). We then carried out Mantel tests based on population averages to exclude the very local distance component. These tests were significant over all diploids (*r*=0.27, *p*=0.002) but not over all tetraploid populations (*r*=0.04, *p*=0.09). However, *E. arctica*, the most widespread tetraploid species, showed a pattern of isolation-by-distance at this level (*r*=0.24, *p*=0.015).

We confirmed a strong relationship between ploidy and latitude (ANOVA *F*_1,190_=18.79, *p*=2.4×10^−5^), with diploids generally limited to lower latitudes (being particularly abundant in southern England, Supporting Figure S2) while tetraploids extend to the very north of Britain. However, there was no significant association between GS and latitude within ploidy levels (treating taxon as a random effect, *t*=0.63, *p*=0.53). We then analysed the data for each of the three widely sampled species individually using linear models (Figure **2c**). There was a non-significant trend for the diploid *E. anglica* (slope=0.013pg/(degree latitude), *F*_1,9_=4.23, *p*=0.07, *r*^2^=0.24). Of the tetraploids, *E. arctica* did show that GS increases significantly with latitude (slope=0.013 pg/(degree latitude), *F*_1,16_=9.36, *p*=0.008, *r*^2^=0.31), whereas *E. micrantha* did not (*F*_1,7_=0.34, *p*=0.577).

### Variation in genomic repeat content

To investigate the nature of the PAVs underpinning GS variation, we analysed the genomic repeat content from whole genome sequencing data in 31 samples using the RE pipeline. RE’s output includes a set of annotated repeat clusters, representing individual repeat types. Our samples included *B. alpina* (Orobanchaceae), 29 British *Euphrasia* samples (six diploids and 23 tetraploids), and one Austrian diploid (Supporting Information Table **S2**). Overall, 69.9% of all *Euphrasia* reads analysed were identified as derived from repetitive DNA (i.e. they formed repeat clusters with genome proportions > 0.01%). The average genomic repeat contents of diploid and tetraploid *Euphrasia* samples differed, being 71.4% and 69.1%, respectively (*F*_1,28_=8.14, *p*=0.008). The repeat content for *B. alpina* was only 42.4%, which is an under-estimate because repeats private to the species may have failed to form individual clusters given our sampling design and cut-off threshold.

The most abundant repeat family, ranging from 25% in *E. anglica* (AN1) to 30% in *E. cuspidata* (CU), was Angela, a type of Ty1/Copia long terminal repeat retrotransposon (LTR), which is typically c. 8.5 kbp in length. Overall, all types of Ty1/Copia elements identified accounted for 30-39% of each *Euphrasia* genome, while Ty3/Gypsy elements typically occupied just 3-6% of the genome (Supplementary Information Table S2).

To assess how well genomic repeat profiles in samples from different populations correspond with species identity based on morphology, we used two unsupervised machine learning techniques: hierarchical clustering and principal component analysis (PCA). We focussed our analyses on the largest 100 repeat clusters, which together account for approximately 50% of each genome, no matter if diploid or tetraploid. Each smaller repeat cluster had a genomic proportion of < 0.7% in each sample. Hierarchical clustering resulted in a tree that grouped samples largely by ploidy, rather than species identity, with the exception of (i) a sample of the Austrian alpine *E. cuspidata* (CU), a species considered diploid, which grouped as sister to the tetraploids, and (ii) tetraploid *E. arctica* from Cornwall (AR5), which grouped as sister to all other *Euphrasia* samples (Figure **3a**). All species with multiple samples formed mixed branches with other species in this tree. Among the sympatric samples from Fair Isle, *E. micrantha* (MI1-3) clustered separately from *E. arctica* (AR1-3) and *E. foulaensis* (FO1-4), both of which were mixed with other species, similar to previous patterns of clustering from SNP-based analyses (Becher *et al*., 2020).

**Figure 3.**
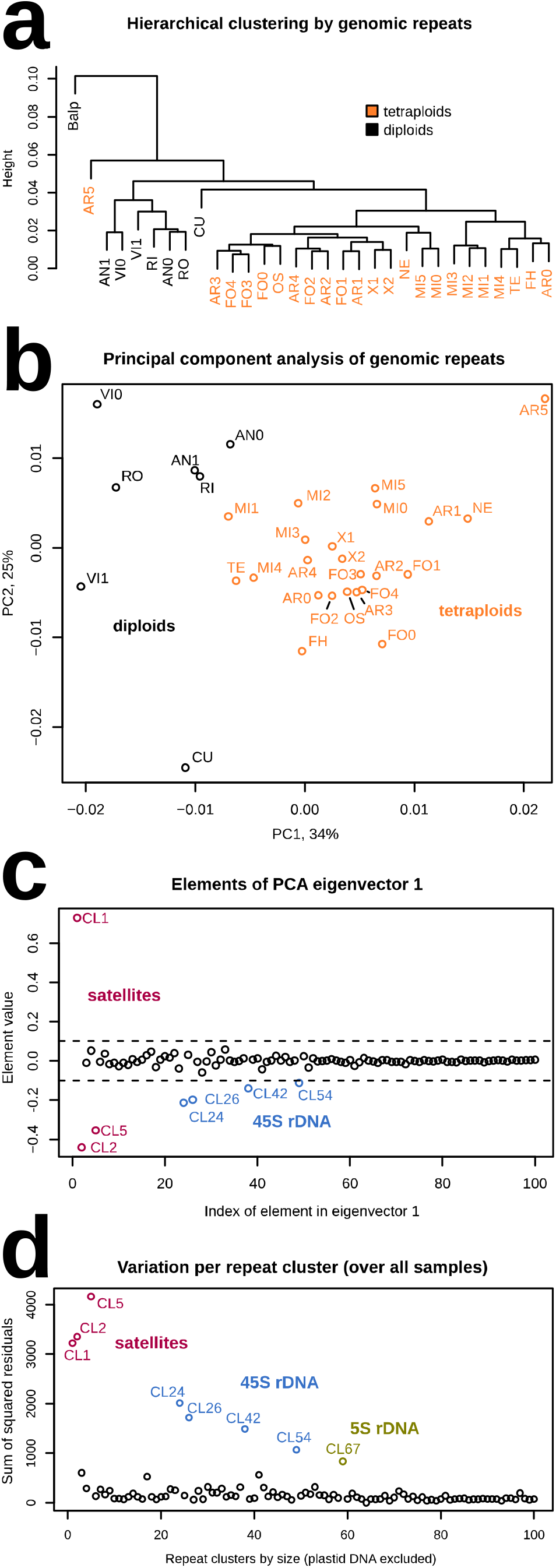
Clustering of *Euphrasia* samples based on genomic repeat content. **a** Hierarchical clustering shows grouping largely by ploidy. **b** A PCA of the relative proportions of the top 100 repeat clusters in 30 samples of *Euphrasia*. Diploids are shown in black and tetraploids in orange. **c** Contribution of each repeat cluster to the first principal component (of panel b). Clusters with negative values are enriched in diploids while those with positive values are enriched in tetraploids. **d** The extent of variation in the genomic proportions of all individuals for each repeat cluster. The codes in **a** and **b** are: Balp-*Bartsia alpina* (outgroup), five diploid species (seven samples): AN-*E. anglica*, CU-*E. cuspidata*, VI-*E. vigursii*, RI-*E. rivularis*, RO-*E. rostkoviana*, seven tetraploid species and two tetraploid hybrids: AR-*E. arctica*, FO-*E. foulaensis*, MI-*E. micrantha*, NE-*E. nemorosa*, FH-*E. fharaidensis*, OS-*E. ostenfeldii*, TE-*E. tetraquetra*, and X-tetraploid hybrids.

PCA without the outgroup *B. alpina* yielded a PC1 that explained 34% of the variance in our repeat data, separating the diploid and tetraploid samples (Figure **3b**), whereas there was no clear separation by species. The samples for some species were spread widely across the plot (e.g. *E. arctica* (AR0-5) and *E. vigursii* (VI0, VI1)), while those of *E. micrantha* (MI0-5) grouped relatively tightly. Although this does not preclude the possibility of species-specific repeat patterns in *Euphrasia*, it is clear that there are no major differences in the relative abundance of the common repeat types between the species. Within the 138 largest repeat clusters, none was species-specific (i.e. present in individuals of only one species). Within the largest 701 clusters, none was diagnostic for a species (i.e., none was present in all samples of one species but absent in all other samples).

To further analyse which repeat clusters separate diploids and tetraploids in the PCA (Figure **3b**), we plotted the elements of eigenvector 1, which correspond to the effect of each repeat cluster on the position of a sample along PC1 (Figure **3c**). Seven repeat clusters have a large effect on PC1, the satellite clusters CL1, CL2 and CL5, and all clusters of the 45S ribosomal DNA (CL24, CL26, CL42, and CL56). Satellite clusters CL1 and CL2 have monomer size peaks of approximately 145 nucleotides as commonly seen in centromeric repeats. In addition, some reads of CL1 and CL2 had paired-end mates in CL22, indicating physical proximity of the repeats within the genome. CL22, in turn, had been annotated as CRM, which is a type of Ty3/Gypsy chromovirus retrotransposon that commonly targets centromeric sequences (Nagaki *et al*., 2003; Neumann *et al*., 2011).

Among all 17 broad repeat types identified by RE (see Supplementary Information Table S2), we found significant differences between ploidy levels for two. Diploid genomes contained higher average proportions of 45S rDNA (4.9%) than tetraploids (2.0%, *F*_1, 28_=20.4, *p*_corr_<0.001), with the genomic proportion ranging from 1.7% to 5.7% in diploids and from 0.8% to 3.4% in tetraploids. Tetraploids contained, on average, more Ty1/copia Ale elements (0.15%) than diploids (0.09%, *F*_1,28_=11.18, *p*_corr_=0.018). While our PCA approach had identified some satellites as highly differentiated in copy number (see above), differences over all satellites were not significant. This is because there was differential enrichment in the ploidy levels for CL1 versus CL2 and CL5 (Figure **3c**). Overall, there is comparatively little differentiation in genomic repeats between the ploidy levels.

We also assessed the variation in repeat content over all samples for each repeat cluster. The eight most variable clusters (i.e. having the biggest differences in repeat proportions between individuals, Figure **3d**), are all tandem repeats (satellites including rDNA). The first seven are the same repeats that separated the ploidy levels in the PCA. The eighth most variable repeat (CL67), which is variable in both ploidy levels, corresponds to the 5S rDNA.

Of the samples analysed with RE, nine tetraploids were from populations which also had GS estimates obtained in this study. Testing the largest 100, 200, and 1000 repeat clusters for correlations between GS and abundance of individual repeat clusters, and correcting for multiple testing by Bonferroni correction, no repeat cluster showed a significant correlation between its abundance in an individual and the population-average genome size. All evidence from repetitive elements suggests that the GS differences between *Euphrasia* individuals of the same ploidy levels are not due to large changes in the genomic proportion of any one specific repeat.

## Discussion

In this study, we investigated the nature of GS variation across taxonomically complex diploid and tetraploid British *Euphrasia*. We complemented an extensive population survey of GS variation with an analysis of genomic repeat composition from seven diploids and 23 tetraploid *Euphrasia*. Overall, we find notable GS variation between populations of the same species, representing a wide range of genuine intraspecific GS variation. Within ploidy levels there is a continuum of GS variation, though ploidy levels have discrete GS ranges. These differences within and between ploidy levels are not attributable to large copy number changes of an individual DNA repeat, but rather to multiple segregating PAVs. Here, we first discuss the link between GS variation and population dynamics and speciation history, highlighting how GS is shaped by many similar processes as population-level sequence variation. We then consider the landscape of repeat dynamics and the potential association with *Euphrasia* polyploid genome history. Finally, we consider the wider implications of framing GS variation in a population genetic framework.

### Genome size variation mirrors population genetic patterns

*Euphrasia* are renowned as a taxonomically complex group where species are recent in origin and show subtle morphological differences, and taxa readily hybridise in areas of secondary contact (Gussarova *et al*., 2008; Wang *et al*., 2018a). Previous population genetic analysis have shown genetic variation is not clearly partitioned by species (Kolseth & Lönn, 2005; French *et al*., 2008; Becher *et al*., 2020), particularly in widespread co-occurring outcrossers, with only certain taxa, like the moorland selfing species *E. micrantha*, being genetically diverged. Here, we find GS variation mirrors these findings of population genetic structure inferred from molecular data. Our results add doubt to the distinctiveness of species, with taxa clearly not showing distinct GS ranges indicative of reproductive isolation. Moreover, previous findings have reported a considerably higher mean GS of 2.73 pg for five samples of diploid *E. rostkoviana* from Bosnia and Herzegovina (Siljak-Yakovlev *et al*., 2010) compared with our estimates that ranged 0.62 -0.71 pg. This notable discrepancy raises a number of non-mutually exclusive hypotheses: (1) heterogeneous GS variation within currently named species may be a consequence of different taxonomic concepts applied across Europe; (2) lower GS variation within British *Euphrasia* may be a consequence of hybridisation and homogenisation of GS variation in Britain or a distinct polyploid history elsewhere in Europe; (3) identification problems or technical issues may affect previous GS estimates.

The continuous GS distribution across species boundaries within ploidy levels in *Euphrasia* resembles the findings of Hanušová *et al*. (2014) for species of the lycophyte *Diphasiastrum* at allopatric and sympatric sites. These authors concluded that considerable GS variation within species resulted from introgression from other sympatric species. Depending on the sizes and number of segregating PAVs (see Figure **1b** and **c**), hybridisation between divergent populations may homogenise local GS, or introduce GS differences. In our study, three populations from Fair Isle (one *E. foulaensis* x *E. marshallii* and two *E. foulaensis*) located within 5 km of each other show likely signals of introgression. Their GS estimates were more than 10% lower than the mean GS of all tetraploids, including all other Fair Isle samples (Figure **2a**). While these populations might have independently evolved lower GS, it seems more plausible that they share large GS difference variants (such as missing dispensable chromosomes or chromosome regions, Figure **1c**). An explanation of genomic homogenisation in sympatry is in keeping with the growing body of plant research showing gene flow at the early stages of species divergence, or between closely related species (e.g. Strasburg & Rieseberg, 2008; Papadopulos *et al*., 2011; Brandvain *et al*., 2014; Sawangproh *et al*., 2020). Such observations of divergence with gene flow are often coupled with species differences being maintained by a few diverged regions under strong selection maintaining species identities (e.g. Twyford & Friedman, 2015), a possibility we are currently investigating in *Euphrasia*.

Within three of the widespread species that we sampled extensively, we found considerably higher GS variation in the mainly outcrossing *E. anglica* and *E. arctica* than in highly selfing *E. micrantha*. Unlike the outcrossing species, *E. micrantha* shows no increase in GS at higher latitudes. Lower diversity is expected for several reasons in young selfing lineages such as *E. micrantha*. Firstly, selfing reduces the effective population size, resulting in lower genetic variation (Nordborg, 1997), presumably including PAVs. Secondly, the reduced effective rate of crossing over between the chromosomes of a selfing species further reduces the effective population size (Conway *et al*., 1999). Thirdly, selfing species are rarely polymorphic for B chromosomes (Burt & Trivers, 2008), one source of GS variation in the Orobanchaceae, for instance in closely related *Rhinanthus* (Wulff, 1939; Hambler, 1953). Finally, partially selfing species are less likely to acquire GS variants through introgression (e.g. Pajkovic *et al*., 2014). Older highly selfing lineages may, however, have diversified ecologically and become restricted to different habitats, and might evolve GS differences.

### Genome size differences and genomic repeats

We found very low differentiation of genomic repeats between species of British *Euphrasia*, with few species-specific repeats. Consistent with phylogenetic work (Gussarova *et al*., 2008; Wang *et al*., 2018a), there were no examples where all species samples cluster together based on repeat content (Figure **3a**). The fact that species of British *Euphrasia* are closely related and often hybridise, makes lineage-specific large-scale gains or losses of individual repeat groups, as seen in other plants (Piegu *et al*., 2006; Macas *et al*., 2015; McCann *et al*., 2020), an unlikely cause for the observed GS variation. Instead, the observed differences are likely due to changes in numerous different repeats segregating within the *Euphrasia* gene pool. At present, it is hard to tell whether these PAVs comprise numerous individual repeat copies or whether there are (also) larger-scale PAVs like the loss or gain of chromosome fragments as hypothesised in hybridising species of *Anacyclus* (Agudo *et al*., 2019; Vitales *et al*., 2020). The high frequency of hybridisation in *Euphrasia* may lead to increased levels of structural rearrangements due to ectopic recombination, which may be more common between heterozygous genomic repeats (Morgan, 2001).

Between ploidy levels of *Euphrasia*, we found allotetraploids had an 11% lower mean GS compared with the value predicted from doubling the mean GS of diploids. This discrepancy may have originated from genome downsizing, commonly seen during re-diploidisation. It may also be explained by the fusion of two diverged diploid genomes of different size, as seen in allopolyploid *Gossypium* (Hendrix & Stewart, 2005) and *Arabidopsis suecica* (Burns *et al*., 2021). However, the absence of interploidy repeat divergence in *Euphrasia* differs from other allotetraploid systems, where diverged sub-genomes tend to show differences in genomic repeats (Zhao *et al*., 1998; Hawkins *et al*., 2006; Renny-Byfield *et al*., 2015; Dodsworth *et al*., 2020). This lack of repeat differentiation is notable because nuclear k-mer spectra (Becher *et al*., 2020) and rDNA sequences (Wang *et al*., 2018a) suggest considerable sequence divergence between the tetraploid sub-genomes, corresponding to a split of approximately 8 million years (Gussarova *et al*., 2008).

Tandem repeats such as rDNA and other satellite DNAs are generally found to be the fastest evolving fraction of the repeatome, showing divergence in both copy number and sequence between closely related species (e.g. Tek *et al*., 2005; Ambrozová *et al*., 2011; Renny-Byfield *et al*., 2012; Becher *et al*., 2014; Ávila Robledillo *et al*., 2020) and populations (Ananiev *et al*., 1998). We confirmed this in *Euphrasia*, where tandem repeats accounted for the eight repeat clusters with the highest inter-individual variation in genomic abundance (Figure **3d**). While differing across individuals, repeat content did not show any clear signal of divergence between particular species. For example, the comparison between *E. micrantha* and divergent tetraploids such as *E. arctica*, did not reveal a signal of divergence in repeat content. This is surprising not just because of their morphological distinctiveness, but their difference in outcrossing rate, with theory predicting that the copy-number and equilibrium frequency of transposable elements depends on the level of selfing in a population (Morgan, 2001; Dolgin & Charlesworth, 2006). A likely explanation is that the shift to high-selfing in *E. micrantha* is relatively recent compared to the time it takes for the genomic repeat content to reach equilibrium level.

### Evolution of genome size variation

The continuous GS variation within and between *Euphrasia* species, coupled with these differences likely being a product of segregating PAV across the genome, underlines the polygenic nature of GS variation. Regarding GS differences as the result of segregating (i.e. genetic) variants blurs the classic distinction between genotype and nucleotype, where “nucleotype” refers to “conditions of the nucleus that affect the phenotype independently of the informational content of the DNA”, essentially identical to GS (Bennett, 1971, 1977). Because GS has been shown to be correlated with many traits including cell size, stomatal pore size, the duration of cell division, and life-history differences (e.g. Šímová & Herben, 2012; Bilinski *et al*., 2018; Roddy *et al*., 2020), it is plausible the GS is affected indirectly by selection on such traits. There might be additional indirect selection on GS according according to the mutational-hazard hypothesis (e.g. Lynch, 2011), which proposes that large GS may be selected against because there is more opportunity for the accumulation of deleterious mutations.

It follows that individual PAVs may be under different kinds of simultaneous selection, potentially of different directionality. For instance, there might be positive selection on an adaptive insertion, which is simultaneously selected against because it increases GS. Further, because selection at one locus affects regions that are physically linked (i.e. selection at linked sites, Maynard Smith & Haigh, 1974; Charlesworth et al., 1993), the footprint of selection on genome regions is modified by the (effective) rate of crossing over, which varies along genomes and between mating systems.

Research on GS is somewhat decoupled from studies on sequence-based variation in populations. We suggest future research into GS evolution should consider both patterns of total GS and the population processes underlying this variation. In addition to furthering our understanding of intraspecific GS diversity in *Euphrasia* and other plant groups, answers to these questions will also improve our understanding of GS evolution between species and across phylogenies, which starts at the population level.

## Acknowledgements

ADT was supported by NERC research grants NE/L011336/1 and NE/N006739/1. The Royal Botanic Garden Edinburgh (RBGE) is supported by the Scottish Government’s Rural and Environment Science and Analytical Services Division. JP was supported by a Ramón y Cajal Fellowship (RYC-2017-2274) funded by the Ministerio de Ciencia y Tecnología (Gobierno de España). We thank Alistair Godfrey, Andrew Shaw, Anne Haden, C.W. Hurfurt, Chris Miles, David Harris, David Nash, Dot Hall, Elizabeth Sturt, Francis Farrow, Geoffrey Hall, Graeme Coles, Jim Hurley, John Crossley, John Wakely, John and Monika Walton, Margaret Chapman, Paul Kirby, Philip H. Smith, Rosemary Parslow, S.J. Bungard, and Stephanie Miles for providing *Euphrasia* samples; Natacha Frachon, Laura Gallagher and Ross Irvine for care of plants; Edgar Wong for preparing NGS libraries; Fergal Waldron for laboratory assistance; Deborah Charlesworth for extensive comments on an earlier version of the manuscript.

## Author Contributions

- HB analysed the data with input from MRB and ADT
- ADT, CM, HB, and MRB collected samples.
- CM confirmed species identifications.
- ADT and IJL designed the study.
- RFP, JP, and IJL generated the GS data.
- HB and ADT wrote the manuscript.
- All authors read and commented on the manuscript.

## Data Availability

The whole genome-sequencing data are available from the sequence read archive, Bioprojects PRJNA624746 and PRJNA678958. The scripts and data required to replicate our results are available from GitHub, repository: zzzzzzz (to be added upon acceptance).

## SUPPLEMENTAL DATA

**Figure S1.**
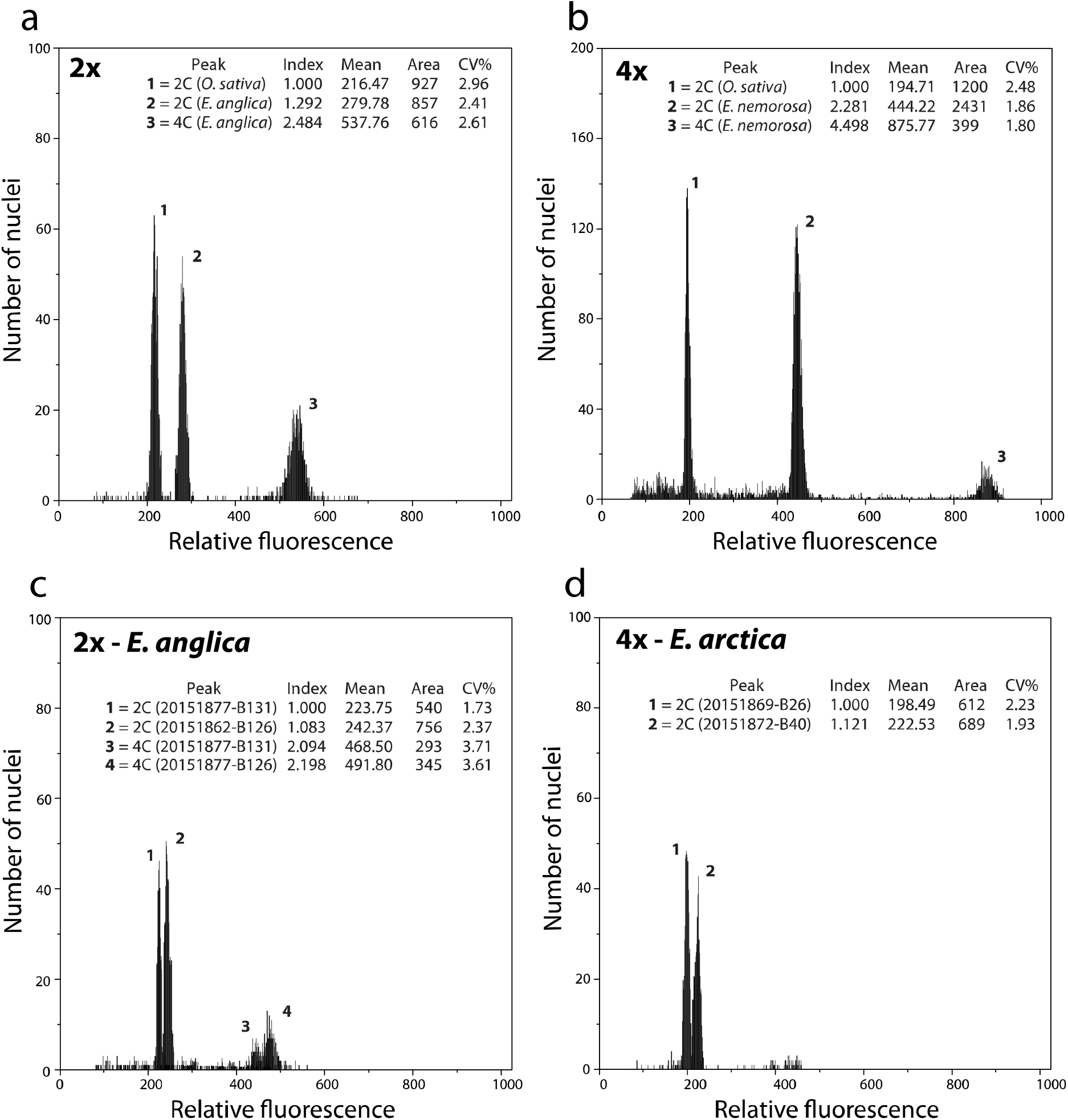
Flow cytometry histograms. A diploid (a) and a tetraploid (b) sample. Intraspecific GS variation in a diploid (c) and a tetraploid (d) species.

**Figure S2.**
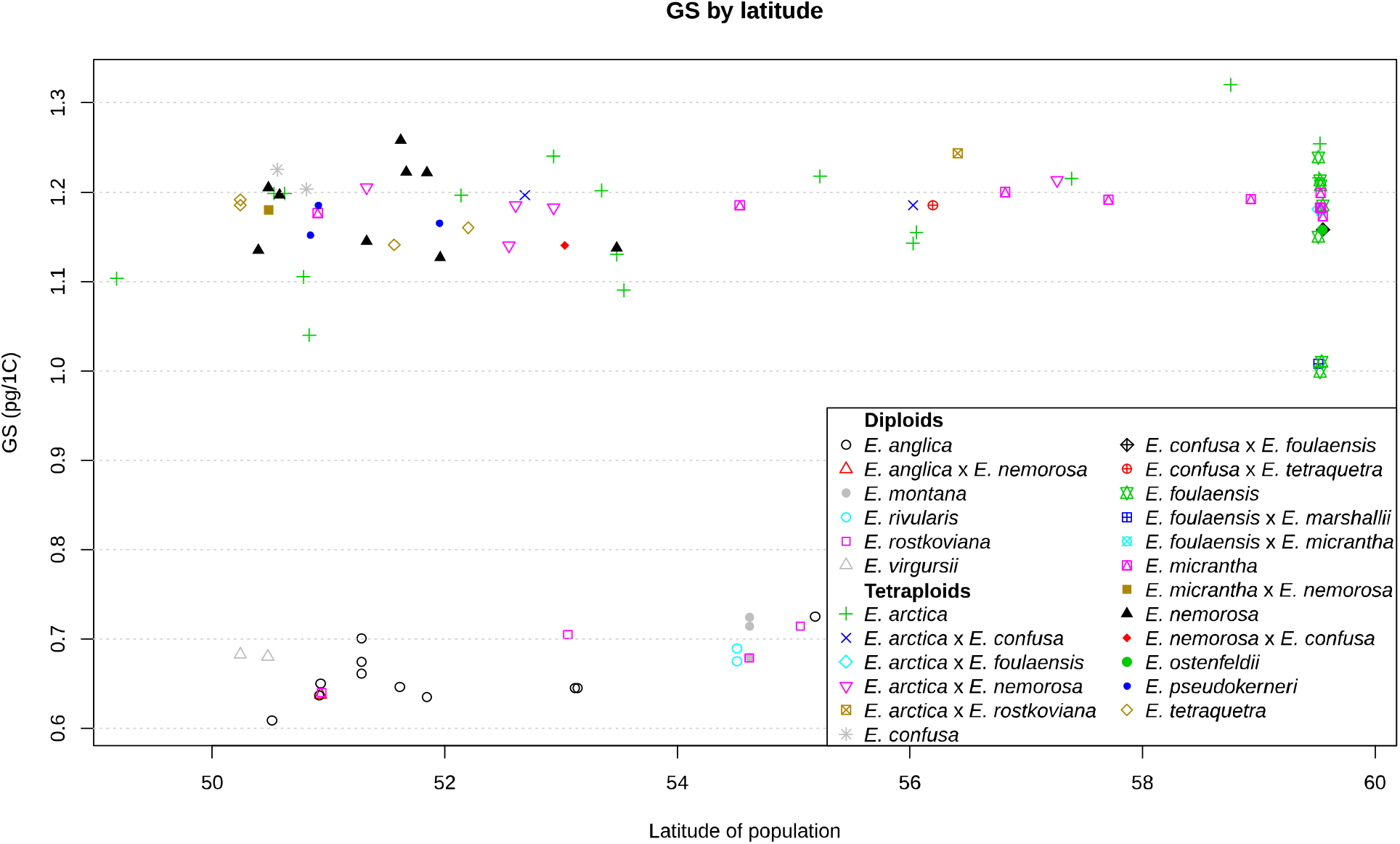
Genome size plotted against latitude.

**Table S1. Sample information and genome size information**

(Submitted separately)

**Table S2. Details of the whole-genome sequencing data sets generated and genomic proportions of repeat types**.

(Submitted separately)

## Notes

### Competing Interest Statement

The authors have declared no competing interest.

